# A freshwater radiation of diplonemids

**DOI:** 10.1101/2020.05.14.095992

**Authors:** Indranil Mukherjee, Michaela M Salcher, Adrian-Ştefan Andrei, Vinicius Silva Kavagutti, Tanja Shabarova, Vesna Grujčić, Markus Haber, Paul Layoun, Yoshikuni Hodoki, Shin-ichi Nakano, Karel Šimek, Rohit Ghai

**Author notes:** Limnological Station, Institute of Plant and Microbial Biology, University of Zurich, Seestrasse 187, 8802, Kilchberg, Switzerland (Adrian-Ştefan Andrei). Authors to whom correspondence should be addressed, Indranil Mukherjee, Rohit Ghai.

## Abstract

Diplonemids are considered marine protists and have been reported among the most abundant and diverse eukaryotes in the world oceans. Recently we detected the presence of freshwater diplonemids in Lake Biwa, Japan. However, their distribution and abundances in freshwater ecosystems remain unknown. We assessed abundance and diversity of diplonemids from several geographically distant deep freshwater lakes of the world by amplicon-sequencing, shotgun metagenomics and CARD-FISH. We found diplonemids in all the studied lakes, albeit with low abundances and diversity. We assembled long 18S rRNA sequences from freshwater diplonemids and showed that they form a new lineage distinct from the diverse marine clades. Freshwater diplonemids are a sister-group to marine isolates from coastal and bay areas, suggesting a recent habitat transition from marine to freshwater habitats. Images of CARD-FISH targeted freshwater diplonemids suggest they feed on bacteria. Our analyses of 18S rRNA sequences retrieved from single cell genomes of marine diplonemids shows they encode multiple rRNA copies that may be very divergent from each other, suggesting that marine diplonemid abundance and diversity both have been overestimated. These results have wider implications on assessing eukaryotic abundances in natural habitats by using amplicon-sequencing alone.

## Introduction

Diplonemids appear as one of the more intriguing findings in amplicon-sequencing surveys of the global oceans microbiota [1]. While the abundance and distribution of these protists were nearly unknown till the last decade, they are now considered (based mainly on metabarcoding approaches) as one of the most abundant and diverse microbial eukaryotic group in the world oceans [2,3]. Marine diplonemid 18S rRNA sequences exhibit high diversity and are the sixth most frequently recovered eukaryotic signatures in the world’s oceans in amplicon-based surveys [3]. They appear to be more abundant in the mesopelagic zone of the oceans [1,4]. These discoveries have ignited interest in understanding the importance and distribution of these protists in other aquatic non-marine ecosystems. Diplonemids are considered marine protists [5] and were till recently never reported from freshwater ecosystems. Our interest in studying the distribution of free-living kinetoplastids (a sister group of diplonemids) from freshwater lakes using kinetoplastid-specific primers accidentally led us to the discovery of freshwater diplonemids in deep freshwater lakes of Japan [6]. Similar to kinetoplastids, diplonemids are generally not targeted by commonly used eukaryotic V4 primers due to their divergent 18S rRNA genes [7,8] and this explains their exclusion from most diversity studies in freshwater environments. This finding suggested that diplonemids are not limited to marine ecosystems and are likely present in other freshwater lakes of the world. To examine the distribution of diplonemids in freshwater habitats, we sampled seven deep freshwater lakes (different depths, both monthly time series and isolated samples) of various trophic statuses from Japan, Czech Republic, and Switzerland. The diversity and abundance of diplonemids were analyzed using PCR based amplicon sequencing, shotgun metagenomic sequencing and CARD-FISH analyses. Our results suggest a broad distribution (albeit with low abundance) of diplonemids in freshwater ecosystems, and the presence of one widespread freshwater clade that appears to be a radiation of known marine isolates.

## Results and Discussions

### Freshwater diplonemid 18S rRNA gene sequences

A single diplonemid sequence (OTU_4) was described recently from the lakes Biwa and Ikeda using 18S rRNA amplicon sequencing [6]. This sequence provided the first intriguing evidence for the existence of freshwater diplonemids, which have been until now considered as marine eukaryotes [5]. However, more conclusive evidence for the presence of these eukaryotes and their distribution in other freshwater habitats is lacking. As the first diplonemid sequence was described from Lake Biwa, in an effort to more clearly examine the depth- or seasonal-related abundance of freshwater diplonemids, we carried out multiple sampling campaigns with different sampling frequencies at this site. The first campaign in Lake Biwa lasted for three months in spring/summer 2016 and samples were collected from three different depths (Table 1, Supplementary Figure S3). The second campaign lasted throughout the year 2017 with a monthly sampling frequency (Table 1, Supplementary Figure S4). A single depth profile was also collected from Lake Ikeda to examine the vertical distribution of diplonemids (Table 1, Supplementary Figure S5). To expand the known distribution of diplonemids outside Japan, we also collected samples from five European lakes (Řimov, Medard, Zurich, Constance and Lugano). A complete list of sampling sites and their locations, depths and analyses performed with the samples from each site is given in Table 1.

**Table 1.** Details of sampling sites

| Lake,<br>country | Trophic<br>status | Surface<br>area (km<sup>2</sup>) | Z max<br>(m) | Sampling depths<br>(m) | Sampling date | Analyses<br>performed |
| --- | --- | --- | --- | --- | --- | --- |
| Biwa, JP | Mesotrophic | 674.0 | 104 | 5, 10, 50 (2016)<br>5, 20, 60 (2017) | Mar-May 2016 (weekly)<br>2017 (monthly) | AS, CF<br>AS, CF |
| Ikeda, JP | Mesotrophic | 11.0 | 200 | 5, 50, 100, 200 | 25.11.2017 | AS, CF |
| Řimov, CZ | Eutrophic | 2.1 | 43 | 0.5, 30 | Apr, Aug, Nov 2016,<br>2017 (monthly) | MG, CF<br>MG, CF |
| Medard, CZ | Oligotrophic | 4.9 | 50 | 5, 20 | 09.07.2019 | MG |
| Constance,<br>CH/DE/AT | Mesotrophic | 536 | 252 | 5, 200 | 03.07.2018 | MG, CF |
| Zurich, CH | Oligo-<br>mesotrophic | 88.7 | 136 | 5, 80 | 11.07.2018<br>13.11.2019 | MG, CF<br>CF |
| Lugano,<br>CH/IT | Eutrophic | 48.7 | 288 | 5, 50 | 02.04.2019<br>05.11.2019 | MG<br>CF |

Abbreviations: JP, Japan; CZ, Czech Republic; CH, Switzerland; DE, Germany; AT, Austria; IT, Italy; AS, 18S amplicon sequencing, MG, shotgun metagenomics; CF, CARD-FISH

Amplicon-sequencing was performed for the sampling campaigns from Lake Biwa and Lake Ikeda (Figure 1A, B, C and Supplementary Figure S3, S4, S5). Relative abundance of diplonemid sequences in these samples was always below 1%. A total of 18 sequences were recovered, somewhat more frequently from the hypolimnion. The highest relative abundance was observed in the hypolimnion of Lake Ikeda (100 m), contributing up to 0.96 % of total sequence reads (Figure 1C). Although rare, they were present in nearly all the seasons in Lake Biwa (Figure 1) in both epilimnion (5 m) and metalimnion (10-20 m) but only occasionally in the hypolimnion (50-60 m). The highest amplicon read abundance in Lake Biwa was observed in the hypolimnion of May (0.5% of total sequences, Figure 1A). Diplonemid sequences were more abundant in the epilimnion and metalimnion during the early part of the spring and as the stratification developed their numbers increased in the hypolimnion (Figure 1A). However, data from the entire year showed the absence of diplonemid sequences in the hypolimnion during summer and autumn (Figure 1B). This discrepancy could be due to the sample collection in different years (spring sampling in 2016 and monthly sampling in 2017) and depths (50 vs. 60 m). However, the low numbers of recovered sequences preclude definitive conclusions regarding seasonal or depth related preferences for diplonemids. Similar results were obtained upon examining the abundance of freshwater diplonemids using an existing shotgun metagenomic timeline from the Řimov reservoir (Figure 1D). Here, diplonemid sequences in the epilimnion were found only during spring or early summer with low abundance (maximum 0.04% of total eukaryotic 18S rRNA reads) (Figure 1D). In contrast to Lake Biwa, diplonemid sequences were found at all seasons in the hypolimnion of the Řimov reservoir with relatively higher proportions (up to 0.6% of total metagenomic reads) (Supplementary Table S2) although these samples were pre-filtered with 5 μm filters, which may have reduced the number of diplonemids. Moreover, the differences in trophic status and sample collection from different depths (Table 1) might be responsible for the observed variation. It is also relevant to mention that dynamics of diplonemids inferred from raw sequence reads is not fully quantitative, as we observed multiple copies of 18S rRNA in single cell genomes from marine diplonemids (Supplementary Table S1).

**Figure 1.**
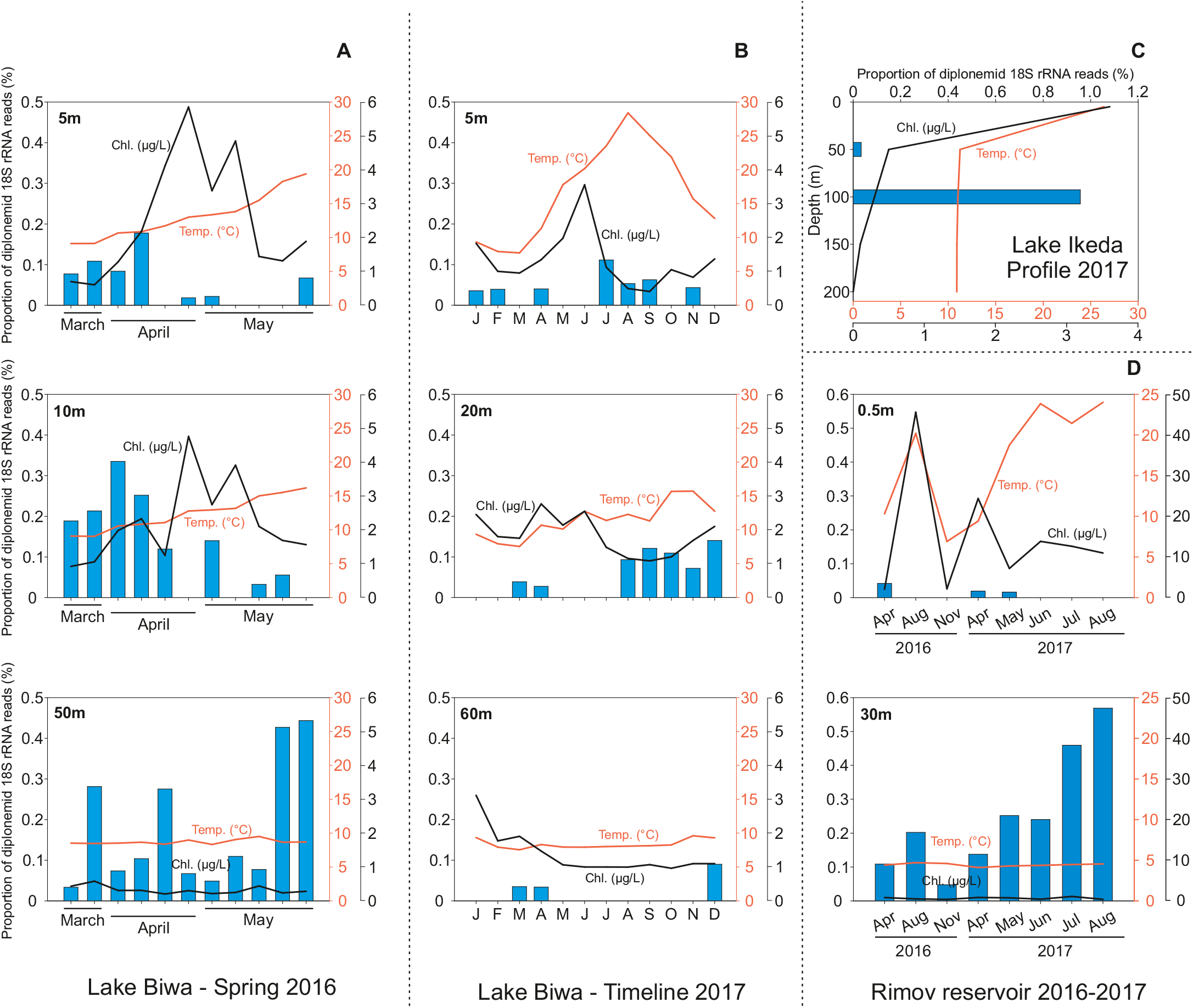
Dynamics of freshwater diplonemids (18S rRNA gene sequence abundance) by A) amplicon sequencing in a weekly spring/summer sampling campaign from Lake Biwa, Japan (depths 5, 10 and 50m), B) amplicon sequencing in a one year sampling campaign from Lake Biwa (depths 5, 20 and 60m) C) shotgun metagenomic time series of the Rimov reservoir, Czech Republic (depths 0.5 and 30m) and D) in a single depth profile from Lake Ikeda (depths 0, 50, 100, 150 and 200m).

To further examine the distribution of diplonemids at other locations, we sampled several European lakes for shotgun metagenomics (Table 1). As the sequences obtained by amplicon-sequencing were short (300 bp), we wondered if it was possible to identify longer assembled 18S rRNA sequences from freshwater diplonemids in assembled metagenomic contigs. Additionally, we also used previously available metagenomic samples from the Řimov reservoir [17] and other published metagenomic datasets from the deep Lake Baikal [18,19]. Multiple long sequences were retrieved from all samples except for Lake Baikal and all of them were closely related to the amplicon-derived sequences (See Figure 2). The longest assembled 18S rRNA sequence was 2029 bp suggesting the less examined potential for assembled metagenomic sequence data for recovery of long eukaryotic 18S rRNA sequences.

**Figure 2.**
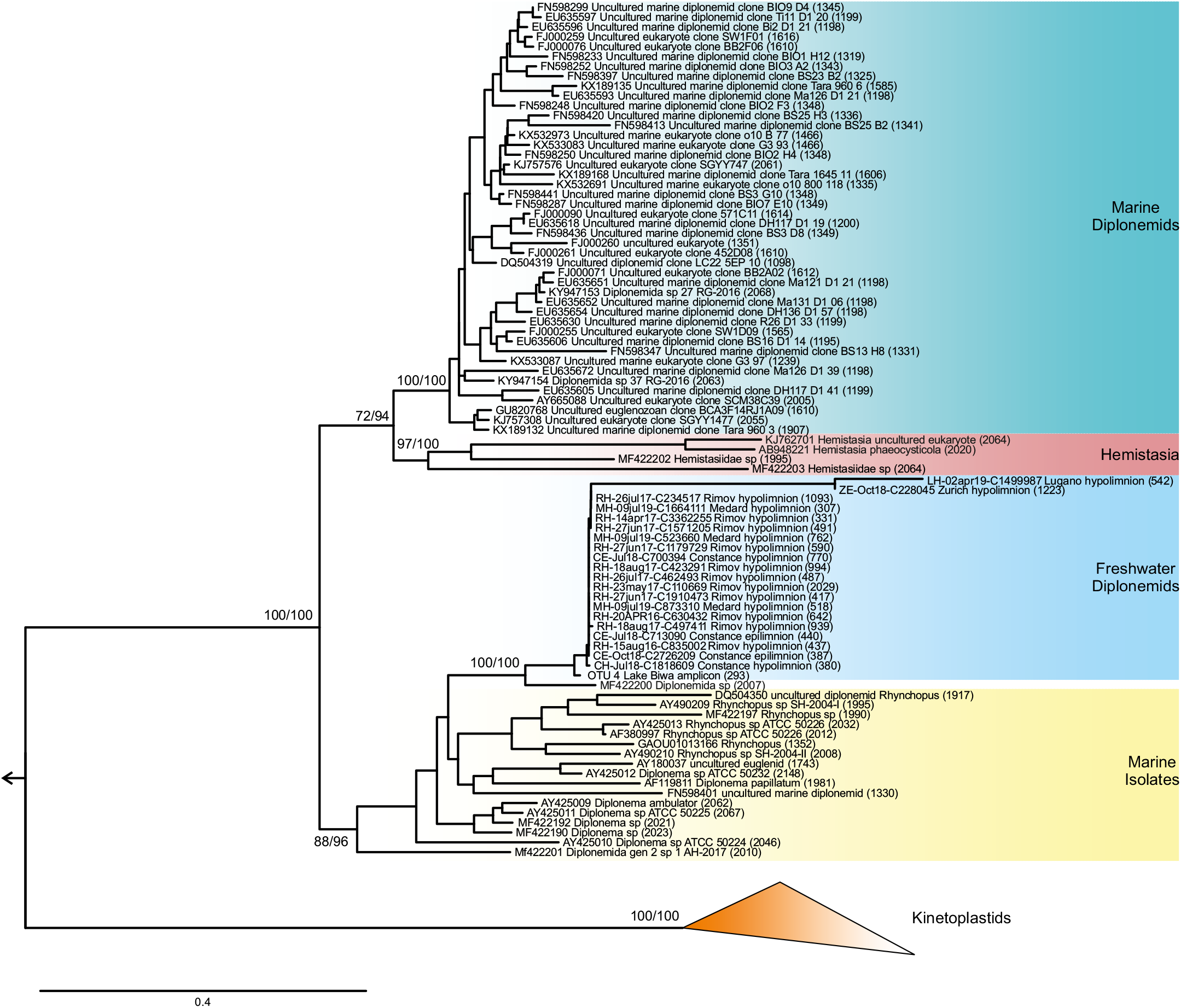
Maximum-likelihood tree of 18S rRNAof diplonemids. Marine diplonemids, *Hemistasia,* freshwater diplonemids, sequences from isolates and kinteoplastids are shown. Other euglenids (e.g. *EUglena, Phacus*) were used as outgroup (not shown and indicated by an arrow). Ultra-fast bootstrap values and SH branch supports are indicated for selected nodes. Scale bar indicates the number of substitutions per site.

Long sequences (>300 bp) of diplonemids from the metagenomic datasets allowed us to construct a robust maximum-likelihood phylogenetic tree including all closely related sequences of diplonemids from public databases (Figure 2). We observed that the sequences recovered from geographically distant freshwater lakes were highly similar and appeared to be a sister clade of several marine isolates. High similarity of diplonemid sequences from freshwater lakes of the world reiterates what has been suggested already [32] that geographic distance does not restrict the distribution of protists in freshwater lakes around the world. The freshwater sequences formed a separate freshwater cluster that was separated from the marine cultured and environmental sequences with high bootstrap values, suggesting that freshwater diplonemids diverged from the marine ones and are phylogenetically different. The highly congruent freshwater clade provides conclusive evidence for an evolutionary radiation of diplonemids in freshwater ecosystems. Intriguingly, a single diplonema isolate (accession MF422200) that was obtained from Tokyo Bay also appeared to be part of this freshwater clade. This might suggest that even though this isolate was obtained in the bay of Tokyo (Supplementary Table S3), it is possible that it is actually derived from the freshwater inflow into the bay. Moreover, the freshwater cluster is closely related to the marine isolates (Figure 2) and majority of those isolates were from bay, estuarine and near-shore regions (Supplementary Table S3) further suggesting a habitat transition of diplonemids from marine to freshwaters.

### Occurrence, morphology, and potential feeding mode of diplonemids

We conducted CARD-FISH with the diplonemid-specific probe ‘Diplo1590R’ [31] targeting all freshwater and marine sequences, to enumerate their abundance in freshwater lakes. However, the number of CARD-FISH stained cells was too low to confidently assess their abundance and dynamics in all the studied lakes. Their abundance was low even during spring, when 18S rRNA read numbers were relatively high. In total, we detected 22 CARD-FISH stained cells from all lakes, again pointing at their rarity in freshwater habitats. However, images of these cells provide some insights about cell size, morphology and potential trophic strategies. All detected diplonemids were ovoid cells in the range of 5-10 μm in length (Figure 3, Supplementary Figure S6). Thus, the freshwater diplonemids observed in the present study are smaller when compared to the cultured marine diplonemids (which are around 20 μm in size), and also the ones observed from marine waters by single cell sorting, which are much diverse in size (7-20 μm) and shape [20].

**Figure 3.**
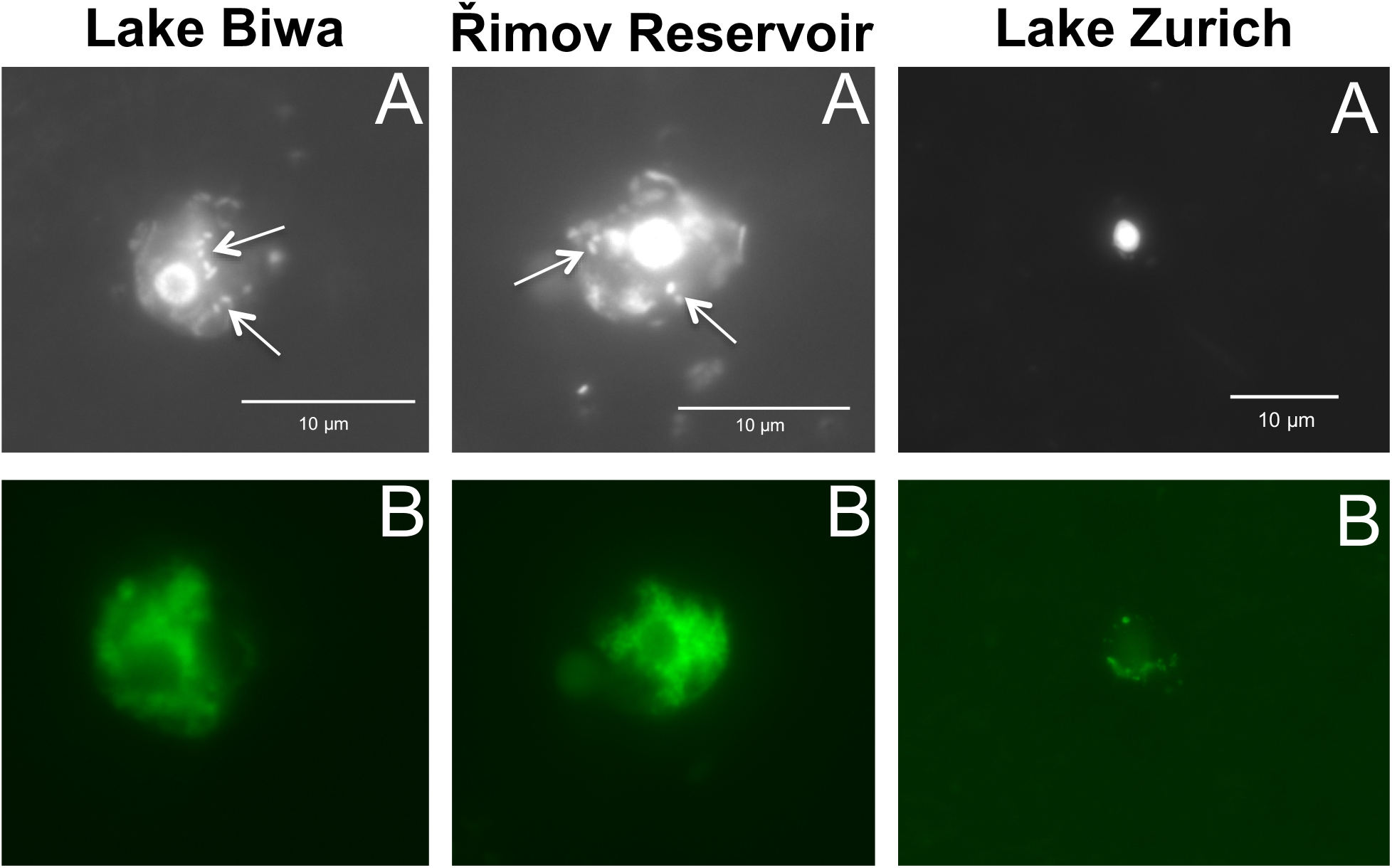
Freshwater diplonemids under fluorescent microscope. A - DAPI, B - FISH, bacteria inside the cells are shown by arrows.

There is little information on the feeding behaviour of diplonemids apart from the consensus that they are heterotrophic. Some studies have also reported diplonemids as mainly benthic dwellers feeding by phagotrophy on bacteria, osmotrophy or by detrivory [33,34]. Protists of size ranges observed here are also known to be efficient grazers of bacteria in aquatic ecosystems [35–39]. All CARD-FISH positive cells observed here also appeared to contain several bacteria (Figure 3). However, it is impossible to conclude from these images whether or not these microbes are inside the cells (suggesting phagotrophic behavior or symbiosis) or outside (epibionts, commensals, etc.) (Figure 3). Although there is no direct evidence of their feeding behavior from environmental samples, diplonemid sequences were detected from the larger planktonic fractions (180-2000 μm) in TARA Ocean data and suggested to possess a parasitic or symbiotic lifestyle [40]. Moreover, a recent study reported the presence of endosymbiotic bacteria inside the cells of some cultured diplonemids [41], which might mimic bacterivory. Direct evidence using FLB (fluorescently labelled bacteria) uptake experiments [42,43] are necessary to confirm their phagotrophy on bacteria.

### Copy number of 18S rRNA genes in diplonemids

Diplonemids have divergent rRNA genes and due to the large number of mismatches, they are generally not targeted in diversity studies using universal eukaryotic primers (especially V4 primers) [7,8]). This explains why they remained undetected in a majority of eukaryotic diversity studies in the world oceans until the TARA expedition. TARA strategically used primers targeting the V9 region [3], which is a short region (~150 bp) considered best to capture a large diversity [10]. Although sequence abundance of diplonemids was high in the global oceanic waters, direct quantitative evidence of their actual abundances is scarce. Only a single study from the Atlantic Ocean quantified the abundance of diplonemids from various depths using CARD-FISH and found their abundance to be <1% of total planktonic eukaryotes [31], similar to our observations from freshwaters. In the microscopic analysis we could barely observe CARD-FISH positive cells even in those samples, which had relatively higher sequence numbers assigned to diplonemids. These observations suggest that diplonemids (especially marine ones) possess high copy numbers of divergent 18S rRNA genes, that could contribute to their amplified sequence abundance [44]. In order to test this, we analyzed ten previously available single cell genomes from marine diplonemids [20]. These genomes are very incomplete (most complete being 9%), so the number of actual 18S rRNAs detected in these is likely to be severely underestimated. Even so, we detected multiple 18S rRNA gene sequences in these incomplete genomes (up to 21 sequences >300bp in cell_13, Supplementary Table S4). Moreover, the sequences from the same cell did not cluster with each other in a phylogenetic tree in several cases (e.g. cell-13, cell-27 and cell-4sb), suggesting a high intra-genomic diversity of this frequently used gene for assessing protist diversity [44] (Supplementary Figure S2). As the raw sequence data for the single cell genomes are not available, we used the coverage of the contigs to obtain an estimate of 18S rRNA copy number (Supplementary Table S4). A conservative estimate, using median coverage of an 18S rRNA containing contig vs coverage of the remaining contigs (>3kb) suggests multiple copies of 18S rRNA genes in one single cell (up to 141 copies, Supplementary Table S1 and Supplementary Table S4). Only a single, related isolate genome is available (*Diplonema papillatum* sp.) with an estimated genome completeness of ca. 30% (Supplementary Figure S1). We could detect 6 copies of 18S rRNAs, which appears to be somewhat low [22]. However, we could not estimate the putative number of rRNAs, as coverage information for contigs was not available. Given such wide variations in the number of 18S rRNA copies and sequence divergence in the presently available incomplete genomic data, accurate quantifications of protist diversity or abundance by sequencing data alone are expected to remain a pressing issue until improved genomes are available.

## Conclusions

We describe a clade of freshwater diplonemids from multiple lakes around the world. Freshwater diplonemids are related to cultured marine diplonemids and distinct from the as yet uncultured marine diplonemids. Direct observations using CARD-FISH also suggest that freshwater diplonemids are consistently smaller in size in comparison to those from the marine habitat. Although diplonemid 18S rRNA gene sequences were found to be highly abundant in marine waters [3], a prior quantification of marine diplonemids using CARD-FISH suggests that they are rare components of the marine habitat [31]. It has also been recently suggested that estimates from amplicon sequencing are far from quantitative [44,45]. Taken together with the highly divergent and multiple 18S sequences derived from single diplonemid cells, it appears the actual abundances of marine diplonemids using sequence data might have been overestimated. Direct counts by CARD-FISH using an identical probe that targets both freshwater and marine clades suggest that diplonemid abundances in both marine and freshwater ecosystems are very low. However, amplicon-sequencing results from both habitats are drastically different, while a large diversity of diplonemid sequences have been obtained from the marine habitat [3], very limited diversity was observed in freshwaters. This suggests that freshwater diplonemids are less diverse than their marine counterparts and their close relationship indicates a recent habitat transition from marine to freshwaters. Nevertheless, it is important to note that this contradictory result of diversity and abundance might not be restricted to diplonemids alone and may also extend to other protistan groups as well [7,43,45]. Further studies using single-cells, cultures, genome sequencing and experimental approaches will be necessary for an improved understanding of the ecological role and evolutionary history of these poorly understood groups of protists.

## Materials and Methods

### Sampling

Samples were collected from seven freshwater lakes, in Japan (Biwa-35°12′ N 135°59′ E and Ikeda-31°14′ N 130°33′ E), Czech Republic (Řimov-48°51′ N 14°29′ E, and Medard-50°10′ N 12°35′ E), and Switzerland (Zurich-47°18’N 8°34’E, Lugano-45°59’N 8°58’E, and Constance-47°32’N 9°31’E) (Table 1). All these lakes are >100m deep with the exception of the Řimov reservoir and Medard, which are 43 and 50m deep, respectively. Samples from Lake Biwa, Japan were collected from three depths representing epilimnion, metalimnion and hypolimnion from station Ie-1, a long-term limnological survey station of Kyoto University, Japan. These samples were collected monthly for one year in 2017 (from depths 5, 20 and 60 m) and also from a high frequency weekly sampling, which was conducted for three months during the spring of 2016 (from depths 5, 10 and 50 m) (Table 1). Samples from the Řimov Reservoir, Czech Republic were collected seasonally for two years (2016 and 2017) from two depths (0.5 and 30 m), representing epilimnion and hypolimnion (Table 1) from a station located at the dam (deepest part of the reservoir). For other lakes, samples were collected once or twice during the thermal stratification period from depths representing the epilimnion and hypolimnion from stations located at the deepest part of the lakes. Samples were collected with a 5-liter Niskin sampler (General Oceanics, Miami, USA) or with a Freidinger sampler. Temperature, concentration of chlorophyll *a* and dissolved oxygen was determined with a multiparameter probe (911 plus, Sea Bird Electronics, Inc., USA; or YSI 6600 multiparameter sonde V2, Yellow Springs Instruments, USA) (Figure 1).

### DNA extraction, amplicon sequencing and data processing

Amplicon sequencing was conducted only from samples collected from Biwa and Ikeda lakes. Water from the different sampling depths was pre-filtered just after collection using a 20-μm mesh plankton net to exclude larger organisms. One to two liters were filtered onto 0.8 μm polycarbonate filters (47 mm diameter, Costar) at low vacuum (7.5 cm Hg) and immediately frozen at −30°C until analysis. DNA was extracted with the Power Soil DNA isolation and purification kit (MoBio laboratories, Carlsbad, CA, USA) and quantified using a Nanodrop ND-1000 spectrophotometer (NanoDrop Technologies, Inc., Wilmington, DE, USA). Polymerase chain reactions (PCR) were conducted using universal eukaryotic primers Euk1391f and EukBr to amplify the V9 region of 18S rRNA genes of all eukaryotes [9,10]. PCRs were performed in 25 μl of reaction volume according to the 18S Illumina amplicon protocol of the Earth Microbiome Project (http://www.earthmicrobiome.org/protocols-and-standards/18s/). Amplicons were examined using agarose gel electrophoresis and DNA was purified using magnetic beads. Samples were quantified using Qubit. The samples were pooled to have uniform DNA concentration in each and were sequenced using the Illumina MiSeq platform. The processing and quality control of the sequencing data were conducted using DADA2 [11]. Chimera check was conducted with DADA2 and also with the reference-based chimera searches against the PR2 database [12]. Amplicons with quality score more than Q30 were retained and amplicons were trimmed to have 300 bp using the ‘Filter and Trim’ function of DADA2. A total of 656,427 reads were obtained after processing. Classification of unique sequences was conducted against the SILVA 132 reference database [13]. The unclassified sequences were additionally examined for their closest relatives with BLAST searches against the NCBI NT database [14].

### Filtration and DNA extraction for shotgun metagenomic analysis

Shotgun metagenomic sequencing was conducted for all lakes except for the Japanese ones (Biwa and Ikeda). Water samples (ca. 10 L each) from lakes Řimov (Czech Republic, depths 0.5 m and 30 m), Medard (Czech Republic, depths 5 m and 20 m), Zurich (Switzerland, depths 5 m and 80 m), Constance (Switzerland/Germany, depths 5 m and 200 m) and Lugano (Switzerland/Italy, depths 5 m and 50 m) were progressively filtered through 20 μm, 5 μm, and 0.22 μm polycarbonate or polysulfone membrane filters (Sterlitech, USA). DNA extraction was performed from the 0.22 μm filters (containing the 5–0.22 μm microbial size fraction) using the ZR Soil Microbe DNA MiniPrep kit (Zymo Research, Irvine, CA, USA), following the manufacturer’s instructions.

### Preprocessing of metagenomic datasets

Shotgun metagenome sequencing was performed with Novaseq 6000 (2 × 151 bp) (Novogene, HongKong, China). Low quality bases/reads and adaptor sequences were removed from the raw Illumina reads using the bbmap package [15]. Briefly, the reads were quality trimmed by bbduk.sh (using a Phred quality score of 18). Subsequently, bbduk.sh was used for adapter trimming and identification/removal of possible PhiX and p-Fosil2 contamination. Additional checks (i.e., *de novo* adapter identification with bbmerge.sh) were performed in order to ensure that the datasets meet the quality threshold necessary for assembly. Each metagenomic dataset (i.e. each sample) was assembled independently with MEGAHIT (v1.1.5) [16] using the k-mer sizes: 49, 69, 89, 109, 129, 149, and default settings. Previously available metagenomes and assemblies from Řimov reservoir [17] and Lake Baikal were included in the analysis [18,19].

### Genomic data processing

Assembled sequence data for single cell derived genomes from 10 published marine diplonemids were downloaded [20]. Genome completeness estimates performed by us (and similar to those reported earlier) for these were found to be very low with the best assembly missing 91% of conserved single copy orthologs (as reported by BUSCO [21]). The assembled *Diplonema papillatum* genome [22] was downloaded from NCBI (accession LMZG00000000). Only the assembled genome is available and raw data was not available although it was mentioned to be 21.5Gb. The genome assembly size is ca. 107Mb (shortest contig 1000bp, longest contig 100kb). The authors estimated the genome size to be 175Mb, but no details were provided on how this estimate was obtained [22]. Genome completeness estimates for all available genomes (*Diplonema papillatum* genome and the 10 marine diplonemid SAGs) were obtained using BUSCO [21] with the euglenozoa lineage (euglenozoa_odb10 version 2019-11-21) (busco -m genome -i genome.fna -o output --lineage_dataset euglenozoa_odb10) (Supplementary Figure S1). Eukaryotic small ribosomal subunit sequences were detected in these genomic assemblies using ssu-align [23]. Only those sequences that were >300bp were retained for further analysis. The results are summarized in Supplementary Table S1.

### Retrieval of 18S rRNA sequences from assembled shotgun metagenomic datasets

All diplonemid sequences were extracted from the SILVA database version 138 [13]. These sequences along with the freshwater diplonemid sequence (OTU_4) described previously [6], were used as queries for BLASTN against the metagenomic datasets. Hits with e-value <1e-5, alignment length >200 bp and percentage identity >90% to the queries were extracted. These candidate sequences were aligned to the global SILVA alignment for rRNA genes using the SINA web aligner [24] and subsequently inserted into the guide tree of SILVA SSU database 138 RefNR 99 in ARB [25] using maximum parsimony and only those sequences that affiliated to diplonemids and of length >300 bp were retained.

### Quantification of 18S rRNA sequences of freshwater diplonemids in a shotgun metagenomic time-series from Řimov reservoir

All assembled freshwater diplonemid sequences (from previous section) along with OTU_4 from Lake Biwa [6] were added to the existing SILVA 138 database of 18S sequences. At least half the reads from each metagenomic dataset (minimum 94 million reads, maximum 277 million reads) were scanned by ssu-align to extract potential eukaryotic 18S reads (minimum ca. 65,000, maximum 180,000 found) and these were used as queries against the SILVA138 database supplemented with freshwater diplonemid sequences using mmseqs2 [26]. Metagenomic query sequences matching the freshwater diplonemid sequences with >95% identity, evalue <1e-5 and alignment length >50bp were considered to originate from freshwater diplonemids (Supplementary Table S2).

### 18S rRNA phylogenetic tree

Assembled 18S rRNA sequences identified to be from freshwater diplonemids were aligned to all known clades of Euglenids using PASTA [27] and a maximum likelihood phylogenetic tree (Figure 2) was made using IQ-TREE 2 using automatic model selection (TIM3e+R10 model), ultrafast bootstraps and SH-aLRT tests (-bb 1000 -alrt 1000) [28–30]. Additionally, all 18S sequences (>300 bp) from the single cell diplonemid genomes and the *Diplonema papillatum* genome were aligned to the same set of sequences and a phylogenetic tree (Supplementary Figure S2) was created as described above.

### Catalyzed Reporter Deposition-Fluorescent in situ Hybridization (CARD-FISH)

Samples for CARD-FISH were collected from the same depths as the samples used for DNA extractions (Table 1). Water samples were fixed in a 2% final concentration of formaldehyde (freshly prepared by filtering through 0.2 μm syringe filter) for at least 3-4 hours before filtration. 50 ml of epilimnion/metalimnion and 100 ml of hypolimnion samples were filtered through polycarbonate filters (pore size 0.8 μm, Advantec), rinsed twice with 1X PBS and twice with MilliQ water, air dried and frozen at −20°C until further processing. CARD-FISH was performed according to the method described previously [7]. The filters were hybridized at 35°C for 12 h with a 0.5 μg ml^−1^ probe concentration and a 30% concentration of formamide. The probe targeting diplonemids ‘DiploR1792’ [31] was purchased from Biomer, Germany. Counting was performed using an Olympus BX 53 epifluorescence microscope under 1000 × magnification at blue/UV excitation. Total microbial eukaryotes were counted simultaneously by DAPI staining under UV excitation. Images were taken and processed using a semiautomatic image analysis system (NIS-Elements 3.0, Laboratory Imaging, Prague, Czech Republic).

## Supporting information

Mukherjee_et_al_Supplementary_Tables

## Data Accessibility

The sequence data of Japanese lakes generated from amplicon sequencing were submitted to European Nucleotide Archive (ENA) and are available under the BioProject: PRJEB37597, BioSamples ERS4413153–226. Sequence data for all metagenomes generated in this work are archived at EBI European Nucleotide Archive and can be accessed under the BioProject PRJEB35770, BioSamples ERS4146214-50 (Řimov reservoir), BioProject PRJEB35640, BioSamples ERS4065084-85 (Lake Medard), ERS4065070-71 (Lake Lugano), BioProject PRJEB35770, BioSamples ERS4409418–21 (Lake Constance), BioSamples ERS4409422–23 (Lake Zurich). All 18S rRNA sequences, alignments and trees are available at figshare (https://doi.org/10.6084/m9.figshare.12091749.v1). This study did not generate any code.

## Author’s Contributions

IM and RG conceived the study. IM performed sampling, amplicon analyses and CARD-FISH. RG performed metagenomic and phylogenetic analyses. MMS, A-ŞA and VSK performed sampling and phylogenetic analyses. A-ŞA, TS, VG, MH, PL, YH, SN, KS performed sampling and DNA extractions. IM and RG drafted the manuscript with the input of all other authors.

## Acknowledgements

We are thankful to the captains of the research vessel ‘Hasu’, Y. Goda and T. Akatsuka for their help in sample collection in Lake Biwa. We are also thankful to Shohei Fujinaga for his help in sample collection and data analysis. P. Znachor, P. Rychtecky and J. Nedoma are acknowledged for help in sampling of Řimov Reservoir and Lake Medard. T. Posch, E. Loher and V. Lanta are thanked for help in the sample collection of Lake Zurich. S. Dirren and the crew of the research vessel ‘Kormoran’ are thanked for help in sampling of Lake Constance, and F. Leporelli and V. Lanta for help in sampling of Lake Lugano.

---

**Supplementary Figure S1.**
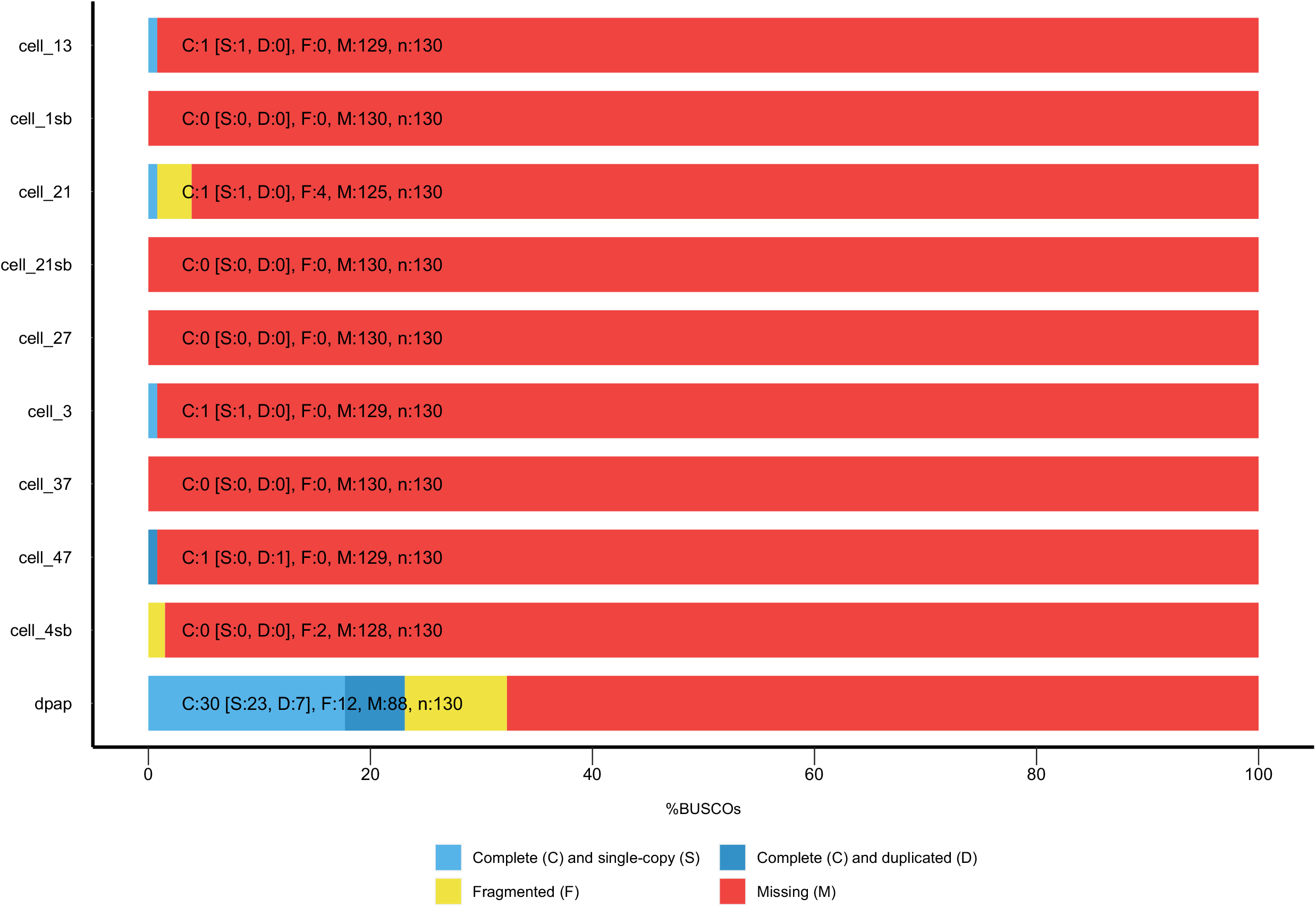
Genome completeness estimation for all available diplonemid genomes using BUSCO (Seppey et al. 2019). Single cell genomes are prefixed with “cell”, Diplonema papillatum isolate genome: dpap.

**Supplementary Figure S2.**
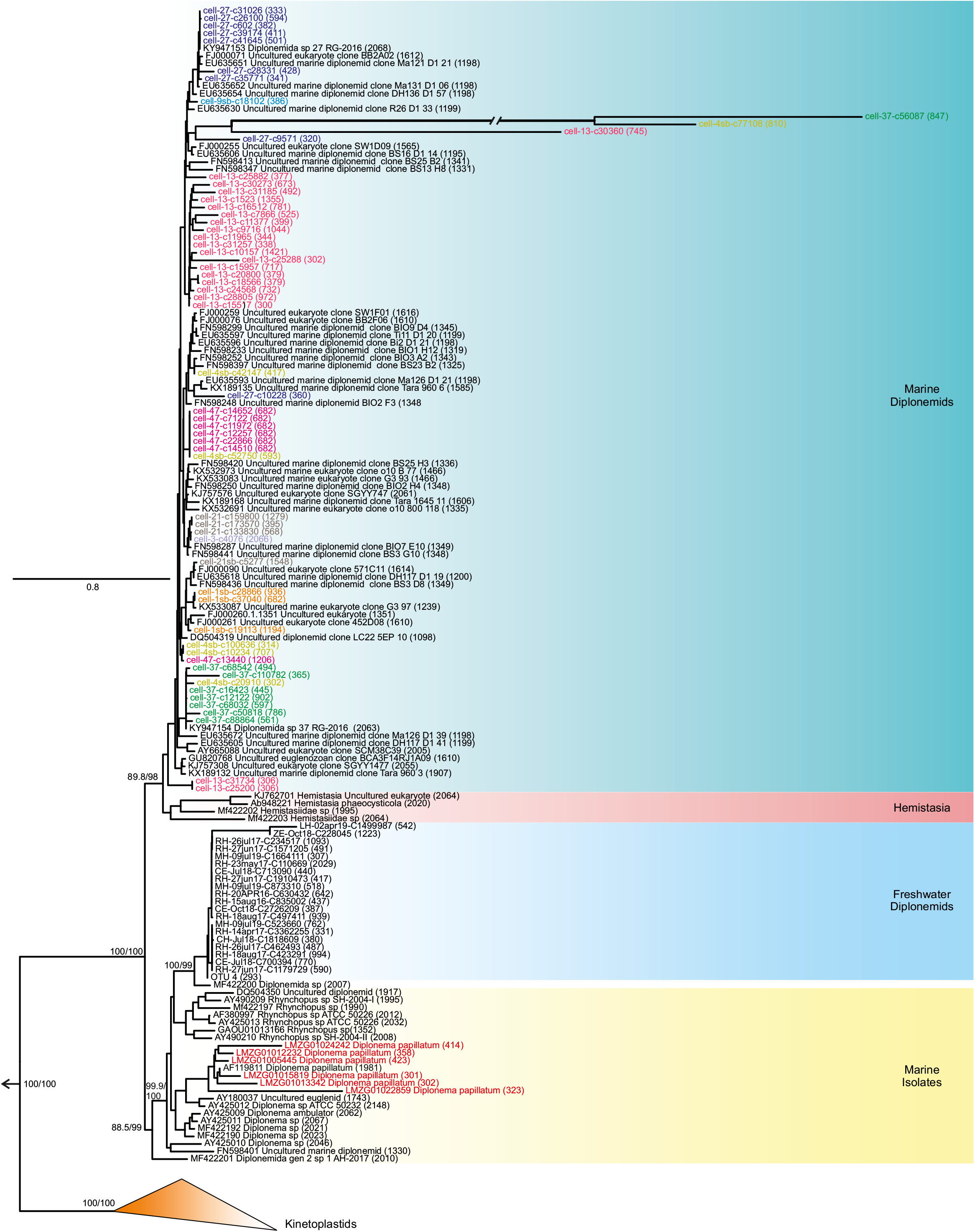
Maximum-likelihood tree of 18S rRNA gene sequences of diplonemids highlighting the copy numbers in single cells. 18S rRNA sequences from the genome of single cells were obtained from public databases and are highlighted with same color. D. papillatum group is highlighted in red in marine isolates. Marine diplonemids, Hemistasia, freshwater diplonemids, sequences from isolates and kinteoplastids are also shown. Other euglenids (e.g. Euglena, Phacus) were used as outgroup (not shown and indicated by an arrow). Ultra-fast bootstrap values and SH branch supports are indicated for selected nodes. The scale bar indicates the number of substitutions per site.

**Supplementary Figure S3.**
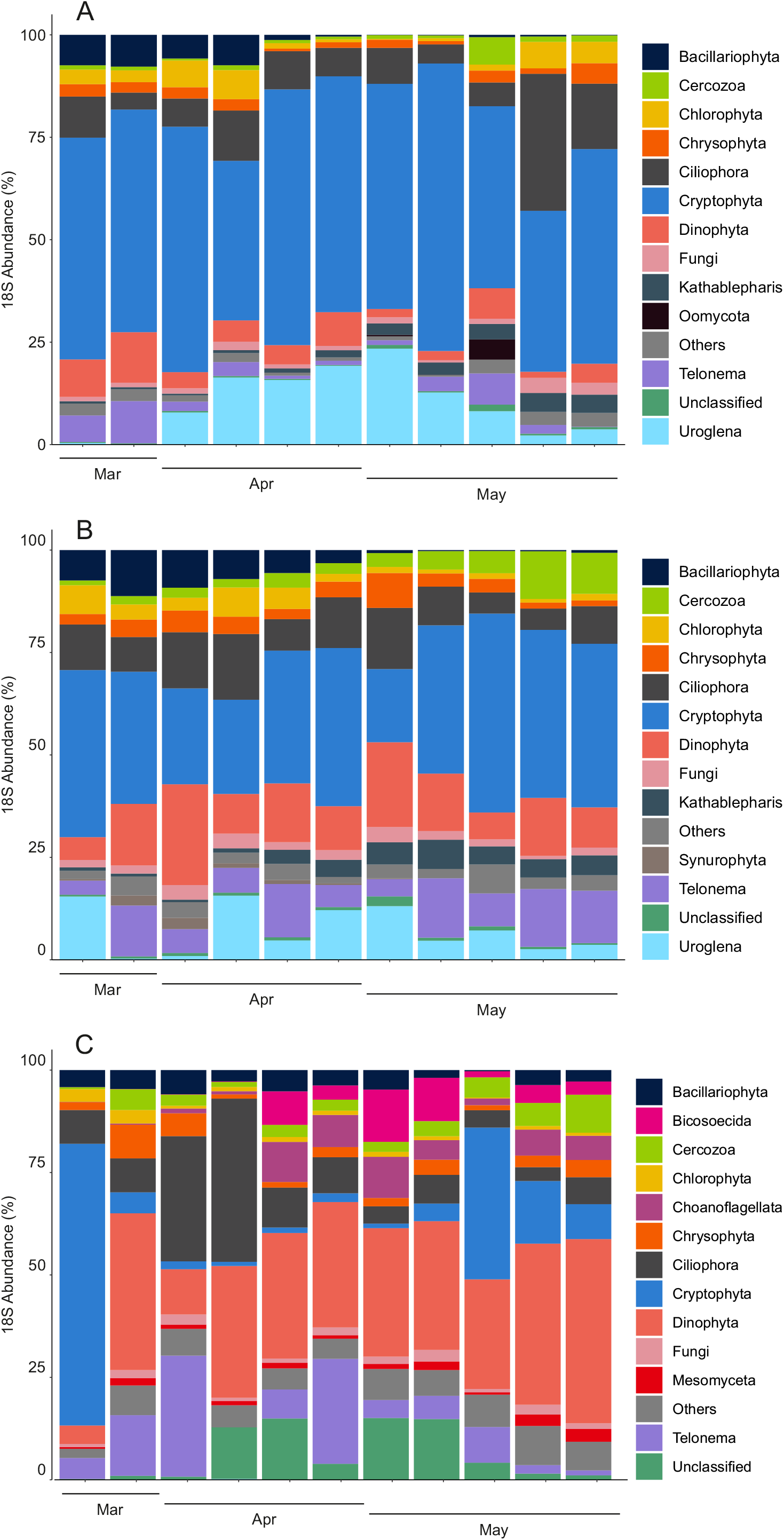
Community composition of microbial eukaryotes in Lake Biwa during weekly spring/summer sampling campaign by 18S amplicon sequencing from A) Epilimnion (5m), B) Metalimnion (10m) and C) Hypolimnion (50m).

**Supplementary Figure S4.**
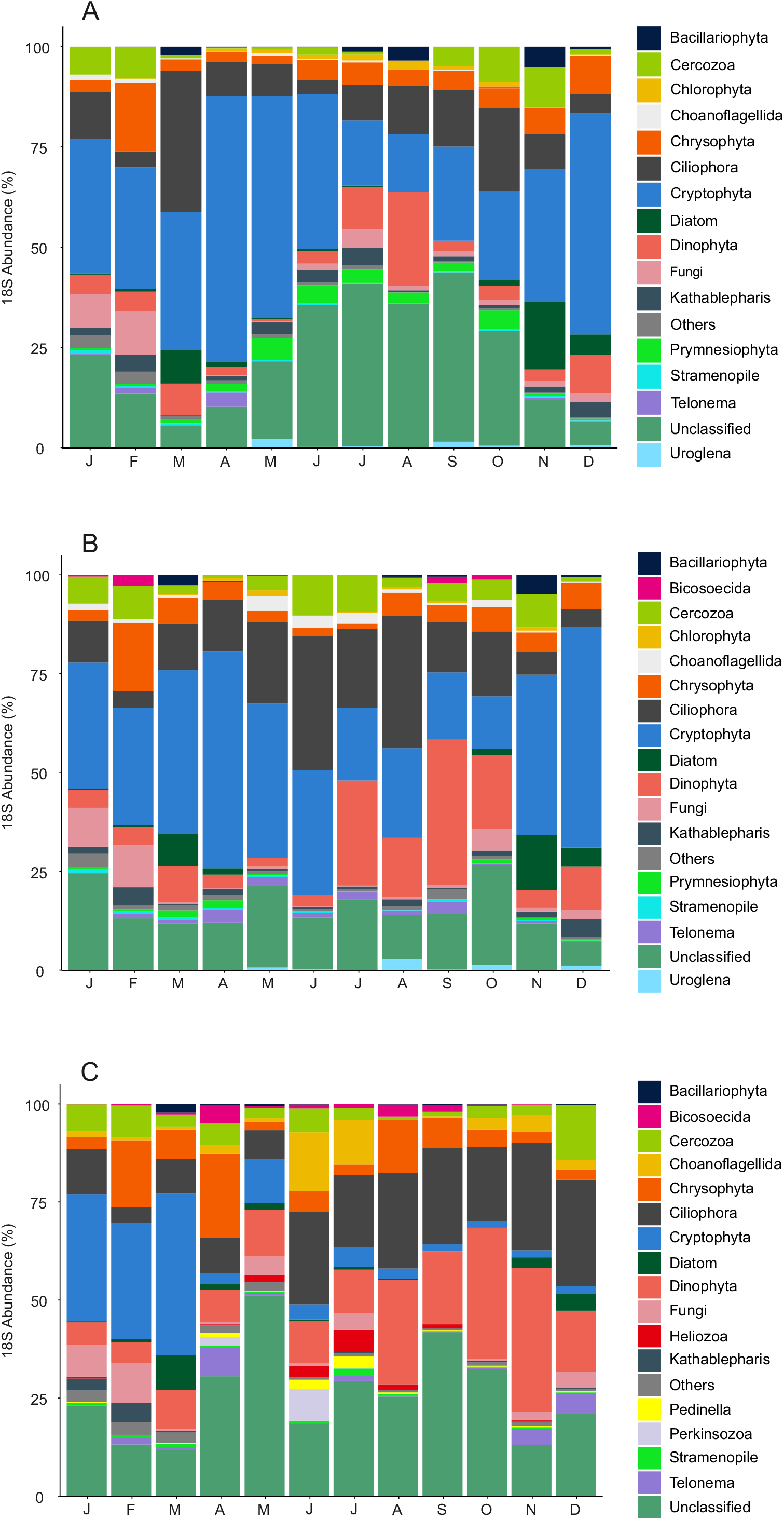
Community composition of microbial eukaryotes in Lake Biwa in 2017 by 18S amplicon sequencing from (A) epilimnion (5 m), (B) metalimnion (20 m) and (C) hypolimnion (60 m)

**Supplementary Figure S5.**
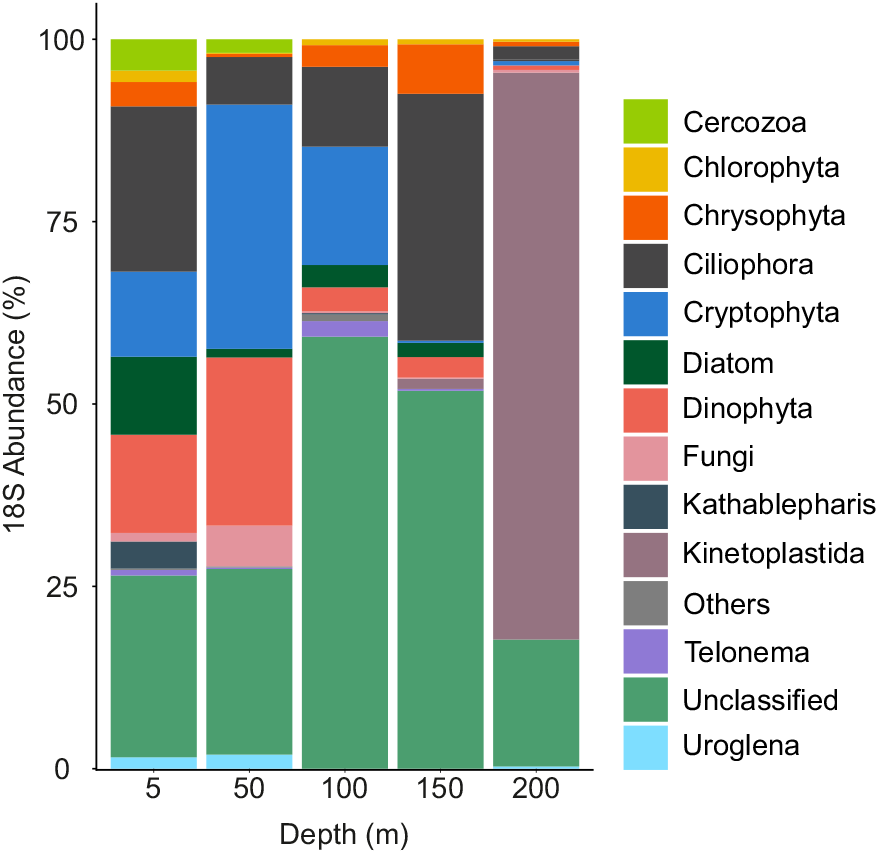
Community composition of microbial eukaryotes in various depths of Lake Ikeda by 18S amplicon sequencing.

**Supplementary Figure S6.**
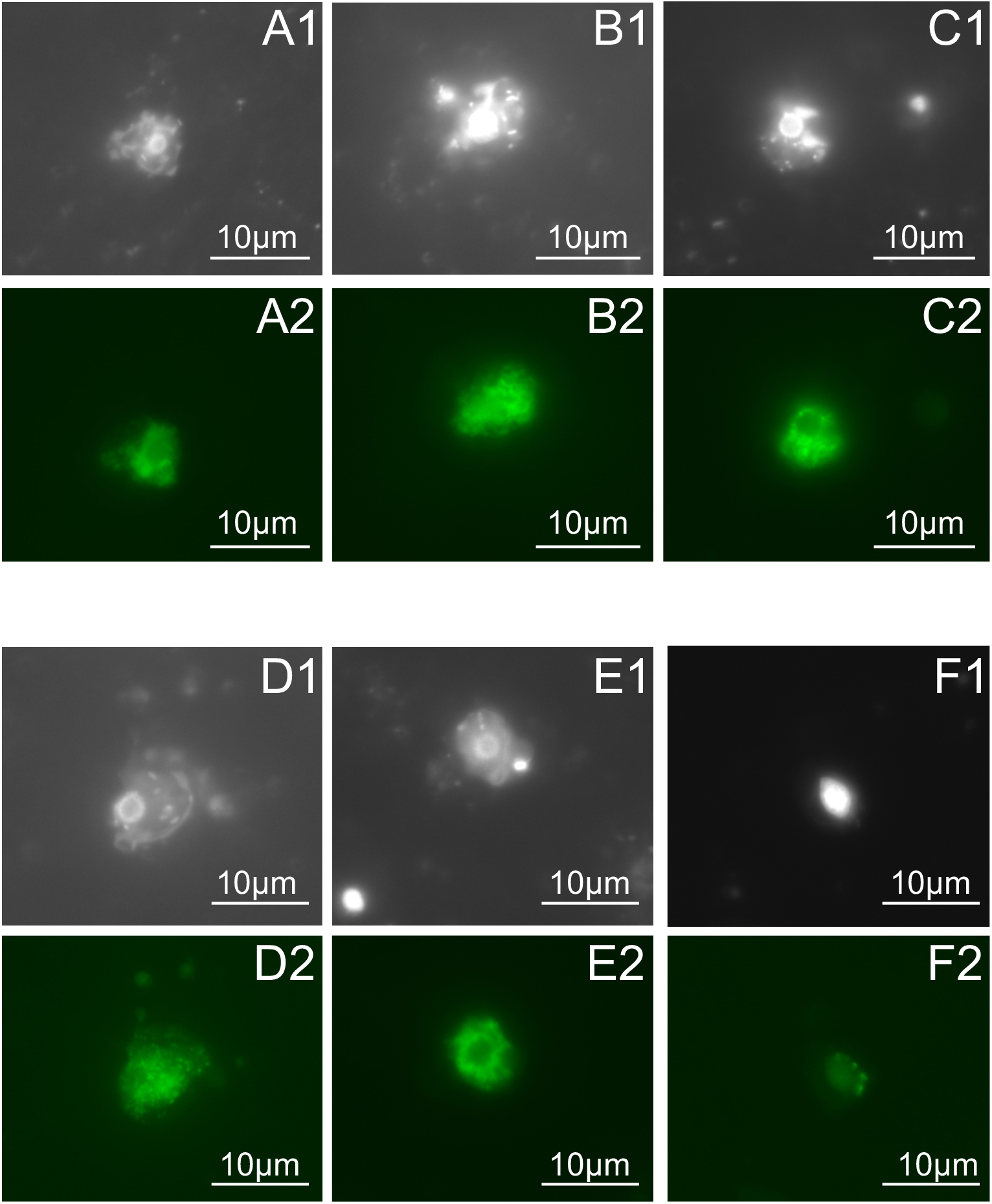
Fluorescent microscopic images of freshwater diplonemids from lake Biwa (A-D), Římov (E) and lake Zurich (F). 1 and 2 refers to the same cells visualized by DAPI (under UV excitation) and CARD-FISH (under blue excitation) respectively.

